# Abundances of transfer RNA modifications and transcriptional levels for tRNA-modifying enzymes are sex-specific in mosquitoes

**DOI:** 10.1101/2021.08.03.454936

**Authors:** Melissa Kelley, Melissa R. Uhran, Cassandra Herbert, George Yoshida, Emmarie Watts, Patrick A. Limbach, Joshua B. Benoit

## Abstract

As carriers of multiple human diseases, understanding the mechanisms behind mosquito reproduction may have implications for remediation strategies. Transfer RNA (tRNA) acts as the adapter molecule of amino acids and are key components in protein synthesis and a critical factor in the function of tRNAs is chemical modifications. Here, we provide an assessment of tRNA modifications between sexes for three mosquito species and examine correlated transcript levels underlying key proteins involved in tRNA modification. Thirty-three tRNA modifications were detected among mosquito species and most of these modifications are higher in females compared to males. Analysis of previous male and female RNAseq datasets indicated a similar increase in tRNA modifying enzymes in females, supporting our observed female enrichment of tRNA modifications. Tissues-specific expressional studies revealed high transcript levels for tRNA modifying enzymes in the ovaries for *Aedes aegypti*, but not male reproductive tissues. These studies suggest that tRNA modifications may be critical to reproduction in mosquitoes, representing a potential novel target for control.

## Introduction

Mosquitoes are vectors for human disease causing millions of deaths each year and compromising 17% of global infectious diseases(World Health Organization, 2020). As such, remediation techniques, such as vector population control are beneficial for preventing disease transmission. Current methods consist of pesticides, genetic modification, and other experimental techniques (Balatsos et al., 2021; Benelli et al., 2016). However, there are limitations to each of these methods and the fear of permanent environmental effects should also be considered (Benelli et al., 2016). Proteomics, transcriptomics, and small RNA sequencing have all been performed in mosquitoes (Camargo et al., 2020; Eng et al., 2018; Gamez et al., 2020; Khan et al., 2005), which has proven critical to understanding mosquito biology. Based on these molecular techniques, males and females exhibit significant differences but there are still gaps in understanding RNA modifications in mosquitoes.

Transfer RNA (tRNA) is responsible for carrying the amino acid to the ribosome based on the codon in messenger RNA (mRNA). As such, the tRNA is critical in protein synthesis. tRNA requires chemical modifications throughout the molecule to ensure stability and alter codon-anticodon interactions (Agris et al., 2007; Edwards et al., 2020; Motorin and Helm, 2010). For example, methylations on the ribose group prevent nuclease cleavage and stabilize the structure of tRNAs to increase thermal tolerance (Hori, 2014). Chemical modifications to tRNA are diverse and range from methylations to more complex hypermodifications (Boccaletto et al., 2018). The hypermodifications are often located on the anticodon and require multi-step synthesis with several enzymes (Boccaletto et al., 2018; Klassen et al., 2016; Noma et al., 2006). Modifications on the anticodon loop may alter the codon preference for a given tRNA. Regardless of type, tRNA modifications have important roles in maintaining translational speed and accuracy.

As tRNA modifications have central roles in translation, they are potential targets for new classes of pesticides and other population control methods. In *Aedes aegypti*, cleavage of tRNA transcripts into tRNA fragments is sex-specific (Eng et al., 2018). As various tRNA modifications have been shown to impact nuclease cleavage of tRNAs (Kawai et al., 1991; Wang et al., 2018), it is possible tRNA modifications are sex-specific as well. However, the tRNA modification status is unknown in mosquito species. In fact, there are few studies on tRNA modifications in insects, particularly in vectors for disease (Koh and Sarin, 2018). In fruit flies, the anticodon hypermodification *N6*-threonyladenosine (t^6^A) was found to be crucial for timely progression into developmental stages and loss of the modification led to smaller larvae likely due to codon-specific defects (Lin et al., 2015; Rojas-Benítez et al., 2017). Still, there has not been a characterization of tRNA modifications or a sex-specific comparison in mosquitoes or other insect species.

Here, we present data that female mosquitoes have a higher abundance of tRNA modifications than their male counterparts. This is supported by tRNA-modifying enzyme transcript levels that are higher in female mosquitoes compared to males. Further, the tRNA modifications detected in *Aedes* and *Anopheles* were sex-specific. Likewise, the relative abundance of many tRNA modifications is higher in females than in males in *Aedes* and *Anopheles*. While *Culex pipiens* exhibited less dramatic differences based on sex, there was one modification elevated in females compared to males and in general, there were slightly higher levels in females. Altogether, female mosquitoes likely utilize chemical modifications to tRNA more abundantly than males, which could underlie factors associated with female reproduction.

## Methods

### Mosquitoes

tRNA modifications will be examined between males and females for *Aedes aegypti, Culex pipiens*, and *Anopheles stephensi*. Larvae were maintained on groundfish food (Tetramin) supplemented with yeast extract (Fisher). All mosquitoes were maintained at 27°C and 80% RH (vapor pressure deficit = 0.71 kPa) under a 15h light: 9h dark cycle with access to water and 10% sucrose ad libitum, unless otherwise mentioned. Mosquitoes were maintained at 27°C and 80% RH under a 15h light: 9h dark cycle with access to water and 10% sucrose ad libitum. Mosquitoes will be 4-5 days old for experiments.

### Identification of tRNA modifying enzymes

To identify possible tRNA modifying enzymes in the mosquito species, peptide sequences of *Homo sapiens* tRNA modifying enzymes were obtained from the UniProt database. Mosquito proteome datasets were acquired from Vectorbase (Giraldo-Calderón et al., 2015). The reference sequences were used to BLAST against the proteome for each mosquito species. A gene was considered to be homologous if the E-value < 1E-29. In **Table S1**, the specific genome for each organism that was surveyed for tRNA modifying enzymes.

### RNA-seq analyses of sex and tissue-specific differences in tRNA modifying enzymes

The SRA experiments used in this study are in **Table S2**. RNA-seq analyses were performed as previously published (Attardo et al., 2019; Scott et al., 2020). First, the adapter sequences were trimmed and assessed for quality using Trimmomatic (version 0.38.0) with default settings (Bolger et al., 2014). Reads were mapped with Sailfish (version 0.10.1.1) under the default settings and generated transcripts per million mapped (TPM) (Patro et al., 2014). Next, DEseq2 (version 2.11.40.6) was utilized to determine differential expression levels following an FDR at 0.05 (Love et al., 2014). To determine sex-specific trends, genes that were two-fold enriched or reduced were considered for pairwise comparisons. The genes enriched in the whole body samples of one sex were considered to be sex-specific. Likewise, tissue analyses were compared to whole body transcriptome levels and the SRA experiments are listed in **Table S2** (Aryan et al., 2020; Honnen et al., 2016; Matthews et al., 2018). These tissue analyses focused on expression levels in relation to the whole body.

### Total RNA and tRNA isolation from female and male mosquitoes

For the RNA isolation protocol, 15 mosquitos were placed in individual bead beater tubes containing 15 beads and 0.5 mL of TRI reagent (Sigma Aldrich). To isolate the total RNA, the samples were inverted and then stored at -80°C until completely frozen. Next, the samples were subjected to bead beating for three cycles of 20 seconds of shaking and 10 seconds of still. Another 0.5 mL of TRI reagent was added and vortexed. Separation was performed by adding 300 µL of acid-phenol-chloroform (Invitrogen), the samples were then vortexed until homogenous and incubated at room temperature for 10 minutes. The samples were centrifuged for 15 minutes at 12,000 rpm at 4 °C. The aqueous phase was collected into a new tube and precipitated with 1.5X volume of isopropyl alcohol (Sigma Aldrich) overnight at -20°C. Total RNA was precipitated by centrifuging for 30 minutes at 12,000 rpm at 4°C. The supernatant was discarded and the RNA pellets were resuspended in 100 µL of 75% ethanol (Thermo Fisher Scientific). The samples were then centrifuged again for 15 minutes at 12,000 rpm at 4°C. The supernatant was discarded again and the pellets were left to dry for 15 minutes. The pellets were then resuspended in 100 µL of sterile water and then stored in the -80°C until tRNA isolation.

For the isolation of tRNAs, total RNA was separated using anion exchange chromatography with a Nucleobond AX100 column (Macherey-Nagel, Duren, Germany). In short, 95 µg of total RNA was resuspended in 100 mM Tris acetate (Research Products International, Mt Prospect, IL) pH 6.3, and 15% v/v ethanol (Thermo Fisher Scientific) solution with pH 7. To equilibrate the column 10 mL of 200 mM NH_4_Cl (Sigma Aldrich), 100 mM Tris acetate pH 6.3, and 15% ethanol solution were added to the column. The total RNA sample was passed through the column three times. Small RNAs are separated in a buffer of 400 mM NH_4_Cl, 100 mM Tris acetate pH 6.3, and 15% ethanol solution. The tRNA fraction was eluted by adding 500 µL of 650 mM NH_4_Cl, 100 mM Tris acetate pH 6.3, and 15% ethanol solution until 12 fractions were collected. The tRNA was precipitated by adding 750 µL of isopropyl alcohol to each fraction, mixed, and stored at -20°Covernight. One biological replicate was defined as tRNA isolated from approximately 200 µg of total RNA and three biological replicates worth of tRNA were collected. Concentrations were assessed via NanoPhotometer (Nanodrop 2000C, Thermo Scientific) and a 1% agarose gel confirmed tRNA presence and purity.

### Nucleoside Analysis of tRNAs to identify modifications

Samples of tRNA were digested into nucleosides using previously reported conditions (Ross et al., 2017). In short, 2 µg of tRNA were denatured by incubating at 95°Cfor 10 min and immediately cooled for 10 min at - 20°C. Next, tRNA was incubated with 1/10 volume of 0.1 M ammonium acetate and nuclease P1 (2 U/µg tRNA, Sigma-Aldrich) at 45°Cfor 2 h. Then 1/10 volume of 1 M ammonium bicarbonate was added. Snake venom phosphodiesterase (1.2×10^−4^ U/µg tRNA, Worthington Biochemical) was added to the mixture to catalyze the formation of individual nucleotides. Phosphates were removed by adding bacterial alkaline phosphatase (0.1 U/µg tRNA, Worthington Biochemical). The mixture was incubated for an additional 2 h at 37°Cand then vacuum dried.

For detection and relative quantification of modifications, nucleosides were resuspended in mobile phase A and separated via reversed-phase liquid chromatography (RP-LC) using a high-strength silica column (Acquity UPLC HSS T3, 1.8 µm, 1.0 mm× 100 mm, Waters). An ultra-high-performance liquid chromatography (UHPLC) system (Vanquish Flex Quaternary, Thermo Fisher Scientific) was used. Mobile phase A was composed of 5.3 mM ammonium acetate in water, pH 4.5, and mobile phase B was composed of a mixture of acetonitrile/water (40:60) with 5.3 mM ammonium acetate, pH 4.5. The flow rate was 100 µL min ^-1^ and the column compartment temperature was set at 30°C. The following gradient was used: 0% B (from 0 to 7.3 min), 2% B at 15.7 min, 3% B at 19.2 min, 5% B at 25.7 min, 25% B at29.5 min, 50% B at 32.3 min, 75% B at 36.4 min (hold for 0.2 min), 99% B at 39.6 min (hold for 7.2 min), then returning to 0% B at 46.9 min.

Next, an Orbitrap Fusion Lumos Tribrid mass spectrometer (Thermo Fisher Scientific) with an H-ESI source (Thermo Fisher Scientific) was used for data obtainment as previously reported (Jora et al., 2018). The analyses were carried out in positive polarity and the settings to obtain the full scan data were a resolution of 120,000, a mass range of 220-900 *m/z*, automatic gain control 7.5 × 10^4^, and injection time of 100 ms. MS/MS fragmentation was carried out with the following collision energy setting for CID from 0 to 42% and the setting for HCD range from 0 to 200 arbitrary units. Other instrumental settings consisted of the following: quadrupole isolation of 1.6 *m/z*; ion funnel radiofrequency level of 30%; sheath gas, auxiliary gas, and sweep gas of 30, 10, and 0 arbitrary units, respectively; ion transfer tube temperature of 299 °C; vaporizer temperature of 144 °C; and spray voltage of 3.5 kV.

Data processing was carried out using Qual Browser in Xcalibur 3.0 (Thermo Fisher Scientific). Three characteristics were used to identify a given nucleoside: retention time (RT), molecular ion *m/z*, and fragment ion *m/z*. Molecular ions *m/z* and fragment ions *m/z* within a mass error of 5 ppm were considered for quantification. Relative abundance was determined by integration of the extracted ion chromatographic peak area which was then normalized to the summation of integrated peak areas of the canonical nucleosides (A, G, C, and U). The relative abundance for three biological replicates of females and males was compared using a Student’s t-test in which a p-value < 0.05 was considered significant.

## Results

### tRNA-modifying expression is higher in female mosquitoes

In general, the expression of tRNA modifying enzymes is significantly higher in females than in males (**Figure 1A**). Moreover, the general trends of expression seem to be held regardless of species. For example, the modifying enzyme FTJS1 is consistently upregulated in females for all three species. In humans, the enzyme FTJS1 methylates many types of tRNAs in the anticodon loop (Nagayoshi et al., 2021). Loss of FTSJ1 reduced the modifications, phenylalanine tRNA levels, and caused slower decoding of Phe codons (Nagayoshi et al., 2021). On the contrary, the tRNA methyltransferase 9B (TRMT9B) is expressed more in males than in females in *C. pipiens* and *A. aegypti* and is the only enzyme that is elevated in males compared to females. TRMT9B methylates wobble position uridines in arginine and glutamic acid tRNAs (Begley et al., 2013).

**Figure 1.**
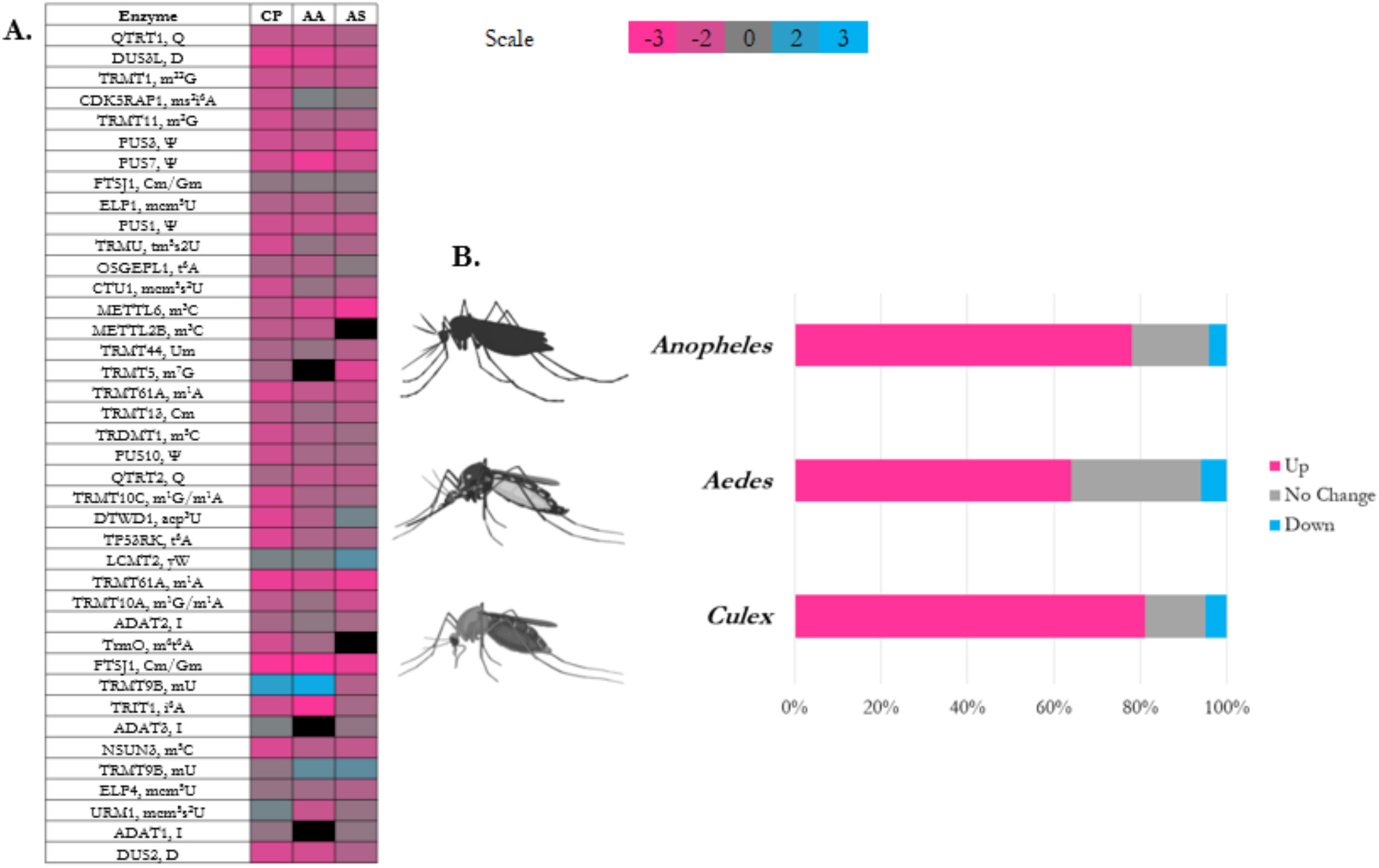
**A**. Homologous tRNA-modifying enzymes identified in *A. aegypti, A. stephensi*, and *C. pipiens*. Values are in log2 and a negative value (pink) indicates the expression is higher in females while a positive value (blue) indicates higher expression in males. **B**. Bar graph of the percentage of differentially expressed transcripts for tRNA-modifying enzymes. The pink bar represents the percentage of tRNA-modifying enzymes with higher transcript levels in females while the blue indicates the percentage higher in males. The gray bar indicates the percentage of tRNA-modifying enzymes that exhibited the same transcript levels in males and females.

A couple of the tRNA-modifying enzyme homologues are for modifications that were not detected in these experiments. For example, in males, TWY4 is upregulated and is a component of the yW biosynthesis pathway. However, the tRNA was absent of yW biosynthesis intermediates (i.e., imG) in any of the samples. Moreover, yW is typically located at position 37 of Phe-tRNA (Guy and Phizicky, 2015). Thus, it is possible yW was not detected due to the low abundance of Phe-tRNA at the conditions the samples were collected. The composition of the tRNA pool likely contributes to the tRNA modification type and abundance detected. Likewise, the male-biased TRMT9B methylates the guanosine (G) at position 34 of Phe-tRNA. The two upregulated enzymes suggest modification of Phe-tRNA is likely required for specific aspects of male mosquito biology. Altogether, differential expression of the tRNA-modifying enzymes suggests there may be alternative modifications present in a sex-biased manner.

### Census of tRNA modifications in mosquitoes

The census of tRNA modifications detected in *A. aegypti, A. stephensi*, and *C. pipiens* is listed in **Table 1**. All four of the canonical nucleosides (A, C, G, U) were detected in each sample as well as Ψ and dihydrouridine (D). Ψ and D are important for the structure of all tRNAs since these modifications are central to the TΨC-loop and D-loop, respectively (Dalluge et al., 1996; Ge and Yu, 2013). Inosine (I) was also detected and is a common wobble modification used to expand decoding capacity. I34 is formed by tRNA-dependent adenosine deaminases (ADATs) and can base pair with C, A, and U on the third position of the codon (Nishikura, 2016; Wolf et al., 2002). As expected, many common ribose and base methylations (e.g., m^5^C, m^3^C, m^3^U, m^5^U, m^6^A, m^1^A, m^1^I, m^2^G, m^1^G, m^7^G, and m^2^_2_G) were detected. In mosquitoes, the 2’-O-methylations detected in tRNA are Ψm, Am, Gm, Cm, and Um. The addition of a methyl group to the canonical nucleoside is common in many kinds of RNA (Boccaletto et al., 2018). In tRNA, methylations are critical for structure. Methylation on the 2’ position of the ribose occurs to prevent nuclease cleavage and provide structural support in tRNA (Kurth and Mochizuki, 2009). Modifications of this type also support structural stability by increasing the melting temperature of tRNA (Ishida et al., 2011).

**Table 1:**
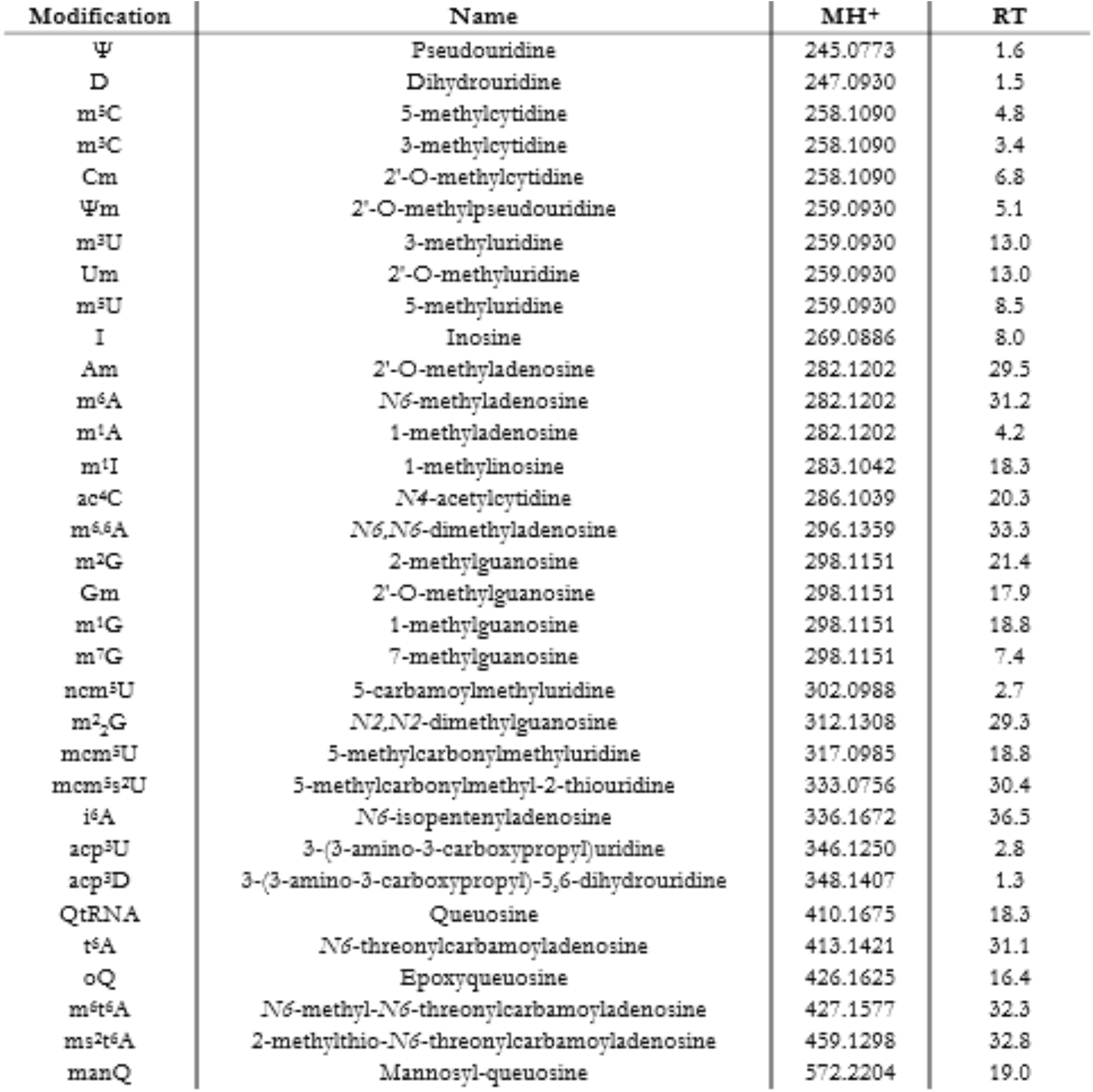
tRNA modifications detected using LC-MS/MS. 33 modifications were detected in all female samples and are listed here. The abbreviation for the modification, the name, the molecular ion m/z, and retention time (RT) are listed here.

One modification is not reported to be present in tRNA and is likely from small contamination of other types of RNA. The modification *N6,N6*-dimethyladeonsine (m^6^2A) has been previously reported in ribosomal RNA of bacteria and eukaryotes (Boccaletto et al., 2018). As a common rRNA modification, presence of m^6^_2_A is likely the result of small ribosomal subunit RNA contamination. However, this modification was detected in low abundance., supporting that this is a minor component. Further, before digestion of the tRNA into nucleosides, samples were visualized on a 1% agarose to ensure there was no rRNA contamination. Thus, if contamination of rRNA is present it is likely minimal.

As for anticodon modifications, several wobble position modifications were detected. Wobble modifications such as these are important in codon interactions and are altered in response to stress in microorganisms (Chan et al., 2012; Fernández-Vázquez et al., 2013). Two of the wobble modifications detected in mosquito tRNA were 5-methylcarbonylmethyluridine (mcm^5^U) and 5-methylcarbonylmethyl-2-thiouridine (mcm^5^s^2^U). The hypermodifications mcm^5^U and mcm^5^s^2^U are performed by the URM and ELP pathways (Rezgui et al., 2013). The type of U34 modification promotes the decoding of certain codons over others. For example, mcm^5^U improves the decoding of G-ending codons (Johansson et al., 2008). In contrast, mcm^5^s^2^U enhances the decoding of A- and G-ending codons (Johansson et al., 2008). In addition, queuosine (Q), epoxyqueuosine (oQ), and mannosyl-queuosine (manQ) are also wobble position modifications. Previously, manQ was thought to always be present with the isomer galactosyl-queuosine (galQ) in eukaryotes (Nishimura, 1983). Q replaces G at position 34 and has only been mapped to four types of tRNAs histidine, aspartic acid, asparagine, and tyrosine (Harada and Nishimura, 1972). The presence of Q on His tRNAs determined the preference of codon (Meier et al., 1985). Altogether, U34 modifications, Q, and its derivatives have roles in codon selection.

The anticodon modifications located at position 37 that were detected in mosquito tRNA were: *N6*-threonylcarbamoyladenosine (t^6^A), *N6*-isopentenyladenosine (i^6^A), 2-methylthio-*N6*-isopentenyladenosine (ms^2^i^6^A), *N6*-methyl-*N6*-threonylcarbamoyladenosine (m^6^t^6^A), and 2-methylthio-*N6*-threonylcarbamoyladenosine (ms^2^t^6^A). Position 37 is located adjacent to the anticodon and serves many functions in regulating translation. For example, t^6^A is universally present across domains of life and stabilizes anticodon interactions (Weissenbach and Grosjean, 1981). Notably, t^6^A is located at position 37 on tRNAs decoding ANN codons. As methionine (Met) tRNAs contain the anticodon CAU, they decode an ANN codon and contain t^6^A (Boccaletto et al., 2018). Met-CAU is also the tRNA that initiates translation by decoding the start codon, AUG. Thus, the presence of t^6^A has been proposed to be a regulator of the initiation of protein synthesis.

### tRNA modifications detected are sex-specific in mosquitoes

Detected modifications in tRNA of *C. pipiens, A. aegypti*, and *A. stephensi* groups have a few specific variations between sexes and species (**Figure 2A**). In *C. pipiens*, sex-biased tRNA modification presence was not observed as all 33 of the modifications were detected in both groups. However, *A. aegpti* and *A. stephensi* demonstrated sex-biased tRNA modifications. Two modifications were absent in *A. aegypti* males that were detected in females. The modifications that were not detected in male *A. aegypti* were oQ and mcm^5^s^2^U. Both of these modifications are present on the anticodon loop and the modification mcm^5^s^2^U is a product of several enzymes and requires multistep synthesis (Björk et al., 2007; Boccaletto et al., 2018). oQ is commonly found in bacteria similar to its final product, Q (Boccaletto et al., 2018; Zallot et al., 2017). Other bacteria-specific modifications (i.e.,2-lysidine or k^2^C) could not be characterized in *A. aegypti* tRNA or the other species.

**Figure 2.**
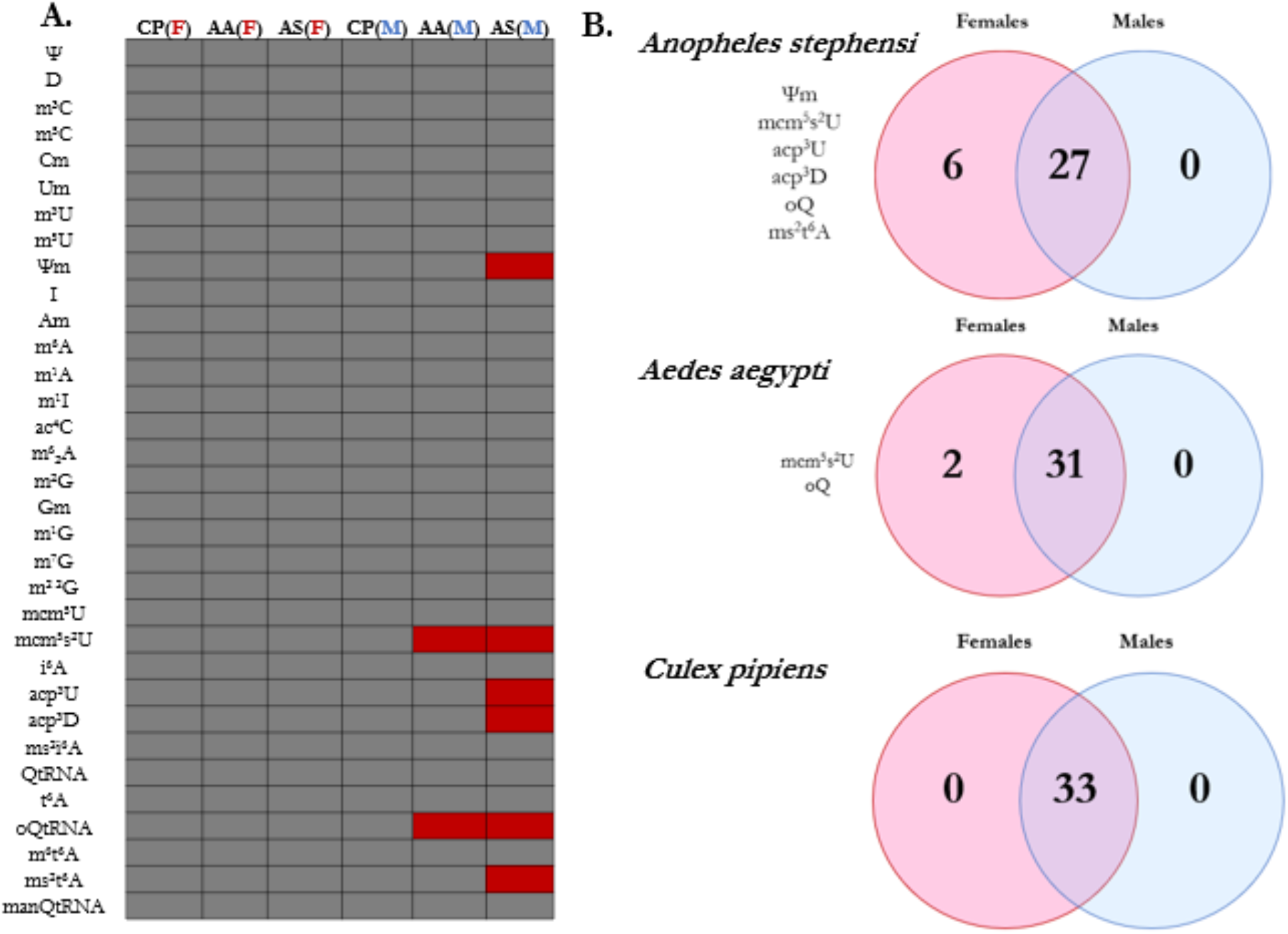
**A**. Modifications detected in tRNA of female and male groups of *C. pipiens, A. aegypti*, and *A. stephensi*. Grey boxes signify the modification was detected while red boxes indicate the modification was absent from that sample. **B**. The Venn diagrams of tRNA modifications detected in each species.

Likewise, oQ and mcm^5^s^2^U were not at detectable levels in *A. stephensi* males. Four other modifications were not detected in *A. stephensi* males: 2’-O-methylpseudouridine (Ψm), 3-(3-amino-3-carboxypropyl)-5,6-uridine (acp^3^U), 3-(3-amino-3-carboxypropyl)-5,6-dihydrouridine (acp^3^D), and ms^2^t^6^A. The modification acp^3^U has been mapped to the variable and D loop of tRNA and the enzymes required for this modification were recently documented in *E. coli* (Meyer et al., 2019; Takakura et al., 2019). While little is known on the function of acp^3^D, the modification acp^3^U has been shown to increase thermal stability and loss of this modification impairs growth in mammals (Takakura et al., 2019). Expectedly, the modification t^6^A was detected across samples and is known to be present in all domains of life (Lorenz et al., 2017). However, t^6^A can be hypermodified to form ms^2^t^6^A or m^6^t^6^A (Boccaletto et al., 2018). The modification ms^2^t^6^A on position 37 has been shown to further stabilize the anticodon interaction with the codon. It is proposed this is accomplished through additional stacking interactions by the ms^2^ group (Durant et al., 2005). Female mosquitoes possess both t^6^A derivatives in tRNA: ms^2^t^6^A and m^6^t^6^A. The modification m^6^t^6^A is proposed to improve the efficiency of the tRNA to read ACC codons in *E. coli (Qian et al*., *1998)*. The presence of both derivatives suggests stabilization in the anticodon loop may be more common in females. However, functional studies of the modifications are necessary to fully understand their role in sex in mosquitoes.

Ultimately, the majority of tRNA modifications are shared regardless of sex and species. Twenty-six modifications were identified in all samples suggesting that the general census of tRNA modifications is relatively conserved in mosquitoes. Furthermore, the detection of modifications is dependent on the composition of the tRNA pool at the time. It is possible modifications present on a single tRNA or tRNA that are in low abundance would not be detected. Thus, the modifications detected in only females may be a product of tRNA transcript abundances in conjunction with modifications.

### tRNA modification abundances are higher in female mosquitoes

In each species, females had a higher abundance of at least one tRNA modification (**Figure 3A**). *C. pipiens* demonstrated the most similar abundances between the sexes with only one modification being higher in females, *N6*-methyladenosine (m^6^A). However, the trend of elevated abundance in females is high for *A. aegypti* and *A. stephensi* which both have heightened levels of at least 13 modifications. Notably, several anticodon modifications were higher in females than in males. For example, m^6^t^6^A and ms^2^t^6^A are both position 37 modifications and were both more abundant in females than males in *A. aegypti* and *A. stephensi*. Another sex-specific anticodon loop modification is Q and manQ. The levels of Q are significantly higher in female tRNA for *A. aegypti* and *A. stephensi*. However, Q abundance is not sex-specific in *C. pipiens* (**Figure 3B**).

**Figure 3.**
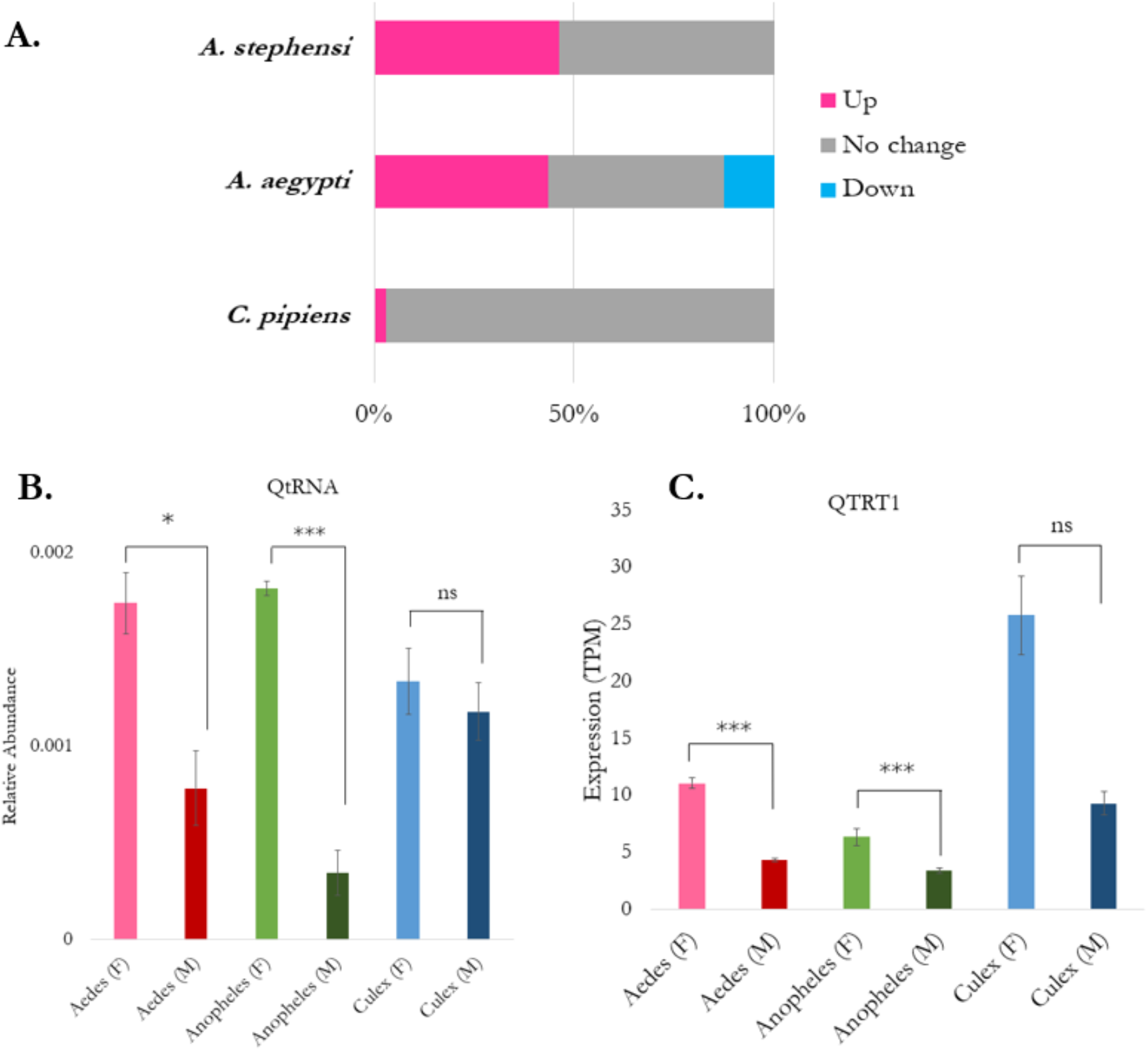
**A**. Percentage of tRNA modifications with sex-specific differences in abundance.The percentage of tRNA modifications exhibiting relative abundance changes as increases in females (pink), no change (gray), or decrease in females (blue). **B**. Relative abundance of the tRNA modification queuosine (QtRNA) in each of the groups. The peak area of QtRNA was normalized with the sum of the canonical peak areas. Significance was considered if p-value < 0.05 between the males and females. **C**. Expression values of the catalytic subunit of the tRNA-modifying enzyme Q transferase, QTRT1, in TPM.

Next, tRNA-modifying enzyme expression differences were compared to tRNA modification abundances. In the case of *A. aegypti* and *A. stephensi*, the catalytic subunit of the enzyme that transfers Q onto tRNAs is Q transferase (QTRT1) follows the sex-specific trend observed in modifications (**Figure 3C**). Likewise, there is not a significant difference in the expression of QTRT1 in *C. pipiens* which correlates with the unchanging modifications levels. We determined general trends in both sets of data to correlate tRNA modifying enzyme expression and modification abundances. To do this, twenty modifications were assessed for the tRNA-modifying enzyme expression in the different sexes. If the modification and enzyme expression is higher in females then the trend was considered to “match”. On the contrary, if the modification levels are not reflected in the enzyme expression, then they are considered not a match (Do not Match). In **Figure 4**, the percentage of tRNA modification abundances that follow the corresponding tRNA-modifying enzyme expression trend is shown for all three species. In *A. stephensi*, the majority of modifications (63%) had abundances that correlated with tRNA-modifying enzyme expression levels. *A. aegypti* tRNA modifications were slightly less in line with the tRNA modifying enzyme expression with only 40% matching expression. Again, *C. pipiens* demonstrates discrepancy in tRNA modification abundance and tRNA-modifying enzyme expression. Only 20% of the tRNA modifications correlate with enzyme expression patterns. Altogether, these differences highlight the complexity of tRNA modification status in mosquitoes, highlighting that expression of modifying enzymes will not likely directly correlate with modification levels.

**Figure 4.**
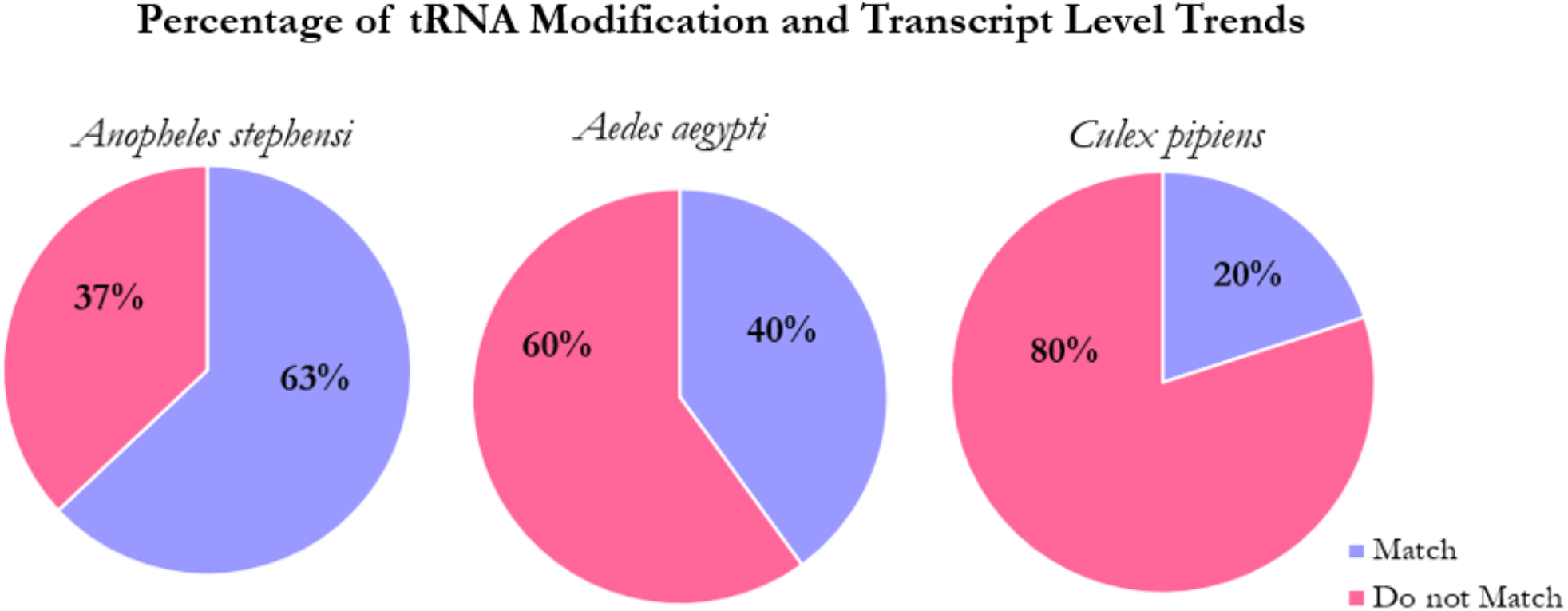
Percentage of tRNA modifications that correlate with tRNA-modifying enzyme transcript levels. Twenty modifications with known tRNA-modifying enzymes were compared to the corresponding transcript levels. A match was defined as a tRNA modification with an abundance that aligned with transcript differences between males and females (i.e, modification and transcript significantly higher in females). The tRNA modification was not considered a match when the tRNA modification abundance did not follow transcript levels (i.e., modification significantly higher but transcript levels were the same).

**Figure 5.**
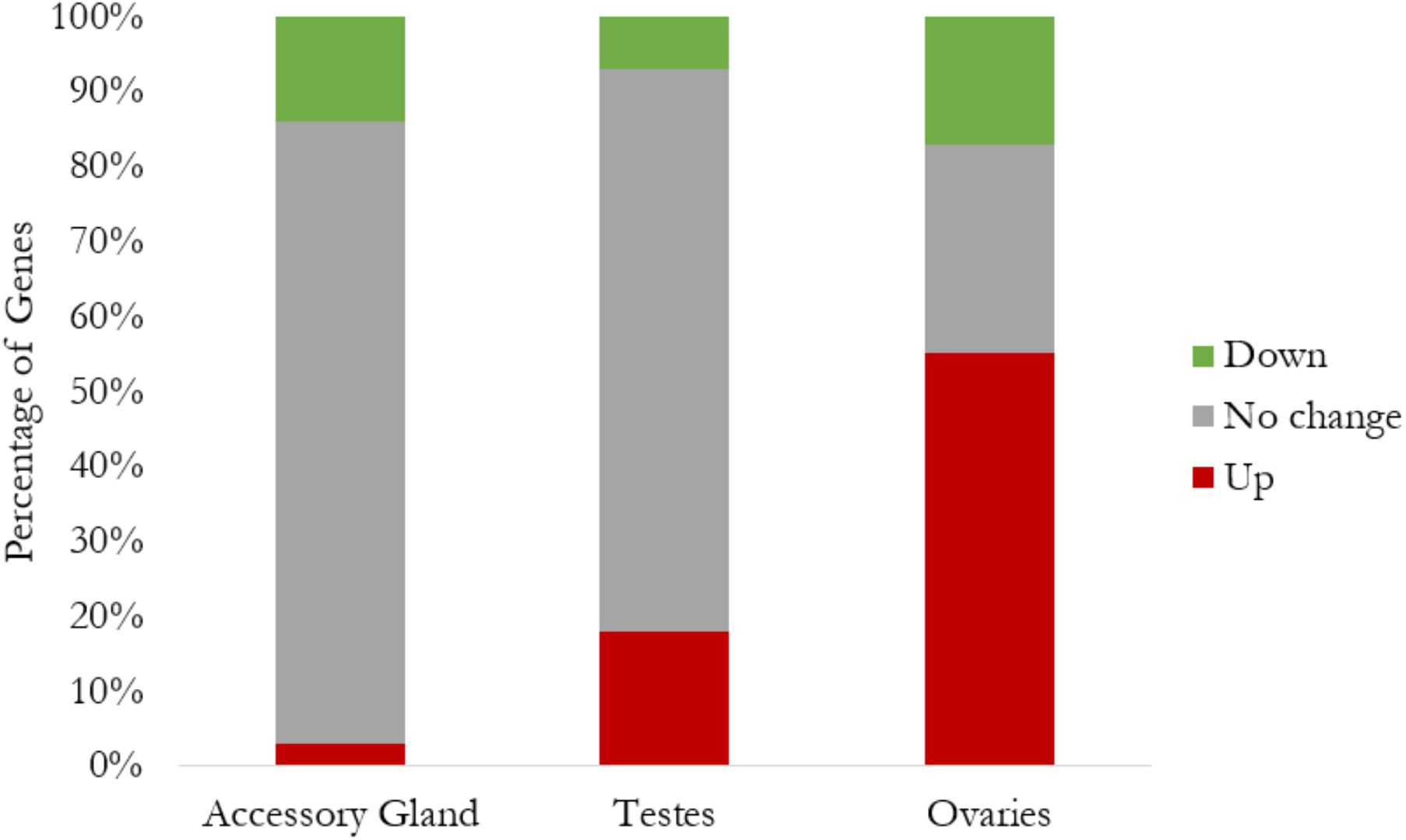
Percentage of tRNA-modifying enzyme gene expression trends in tissues of *A. aegypti* compared to whole body expression. The red bar indicates the percentage of tRNA-modifying enzymes upregulated in the tissue. The gray bar represents the percentage of genes that do not exhibit differential expression between the whole body and tissue. The green bar indicates the percentage of genes that are lower in the tissue than the whole body.

### Reproductive tissues have higher expression of tRNA-modifying enzymes in *A. aegypti*

SRA data sets of isolated tissues in *A. aegypti* were reprocessed. Briefly, expression data were derived from previously published transcriptome experiments on ovaries for female *A. aegypti* and testes and accessory glands in males. The whole body transcriptome experiments were used as a reference. To determine if there is a relationship between tRNA-modifying enzyme expression and reproduction, the percentage of tRNA-modifying gene transcript levels were measured in the accessory glands, testes, and ovaries in *A. aegypti*. Of the three, the ovaries demonstrated the most tRNA modifying enzymes with higher expression when compared to the whole body of a female. Approximately 55% of tRNA-modifying enzymes are induced in the ovaries, suggesting this may be contributing to the overall expression patterns and higher tRNA modifications in females. Enriched transcript levels for tRNA modifying enzymes are higher in testes (20%) and the accessory glands (5%), suggesting a critical role in male reproduction. Regardless, the majority of the tRNA modifying enzymes assessed are differentially expressed in the ovaries of female *A. aegypti*. Ultimately, enrichment in the ovaries implies there may be a biological role of tRNA modifications in the reproduction of female mosquitoes.

In surveying the specific genes enriched in the ovaries, many of the enzymes are for modifications found in every tRNA. For example, *A. aegypti* ovaries had a higher expression of tRNA-dihydrouridine synthase (i.e., DUS2L) which adds the typical D in the D-loop of tRNAs. Likewise, there was a higher expression of tRNA-pseudouridine synthase (i.e., PUS10) which adds Ψ to tRNAs. Enrichment of these enzymes along with many tRNA-specific methyltransferases potentially suggests tRNAs are modified differently in the ovaries than the whole body. However, tRNA modifications do not always reflect expression levels and therefore the tRNA modifications must be assessed for insight on tissue-specific trends.

Other enzymes upregulated in the ovaries were for anticodon loop modifications. Both Q transferase subunits (i.e., QTRT1 and QTRT2) had higher expression in the ovaries than the whole body. Q transferase catalyzes the exchange of guanosine with Q at the wobble position of GUN anticodons (Harada and Nishimura, 1972). Several enzymes that modify position 37 were also enriched in the ovaries. While the enzyme that catalyzes the formation of t^6^A was lower, the enzymes involved in the formation of t^6^A-hypermodifications, m^6^t^6^A and ms^2^t^6^A, exhibited higher expression in the ovaries.

## Discussion

Altogether, we present that female mosquitoes have a higher abundance of tRNA modifications than male mosquitoes. This is supported by enrichment in transcript levels for genes associated with tRNA modifications. This sex-specific impact is more substantial in *A. aegypti* and *A. stephensi* when compared to *C. pipiens*, but even so, this species does show more enrichment. Tissue-specific analyses highlighted that there is high enrichment in the ovaries, suggesting that the increase in modifications and enzyme expression in females may be associated with reproduction. The substantial differences between sexes and enrichment in reproductive organs suggest that tRNA modifications could have critical importance to mosquito biology. Ultimately, the general trends of higher abundance of modified nucleosides and expression in females further suggest tRNA modifications may have roles involved in reproduction.

The tRNA modifications detected in mosquito tRNA are largely shared between the three species. While *A. stephensi* males had the least detectable modifications, the females followed the other species and thirty-three modifications were present. Of the thirty-three modifications characterized, at least nine are known to be located at the wobble position of tRNAs (Boccaletto et al., 2018). As for the other anticodon modifications, six nucleosides were identified that have been previously reported at the position adjacent to the anticodon (Boccaletto et al., 2018). Intriguingly the majority of differences between the males and females were modifications on the anticodon loop. As anticodon loop modifications affect the decoding of codons, the sex-specific differences suggest females have different codon usage than males. Oligonucleotide analysis to map the modifications would confirm the locations of tRNA modifications; however, there are nearly 1,000 tRNA genes in *A. aegypti* which would make differentiating tRNAs a challenge (Chan and Lowe, 2016). Altogether, the census of tRNA modifications is largely the same with 79% of the modifications being shared in all mosquito tRNAs.

A blood meal is required for reproduction in all three of the mosquito species assessed (Attardo et al., 2005; Hansen et al., 2014). However, a blood meal also induces several stresses on the organism including thermal stress (Benoit et al., 2019, 2010; Benoit and Denlinger, 2017). Chemical modifications to tRNA contribute to thermal stability and thermophilic organisms often have higher levels of methylation in tRNAs (Hori et al., 2018; McCloskey et al., 2000). For example, in the thermophile *Thermus thermophilus* unmodified phenylalanine tRNAs had a lower melting temperature than those of modified transcripts (Tomikawa et al., 2010). Furthermore, 2’-O-methylations (Am, Gm, Cm, Um, Ψm) increased the melting temperature of tRNA by more than 20°C in another thermophile (Noon et al., 2003). Thus, tRNA modifications are a strategy to combat elevated temperatures by reinforcing structure and stability. In female mosquitoes, four methylations are in higher abundance in tRNA of *A. aegypti* and *A. stephensi*. In addition, *C. pipiens* females had one modification more abundant in tRNA and it is the methyl modification m^6^A. While additional functional studies are necessary, the higher incidence of methylations in tRNAs of female mosquitoes is potentially in preparation for the thermal stress brought on by a blood meal.

Substantial transcriptome remodeling also occurs in females in the hours following a blood meal. For example, *Culex quinquefasciatus* and other mosquitoes upregulate the egg yolk protein vitellogenin post-blood meal (Isoe and Hagedorn, 2007; Reid et al., 2015). This is accompanied by many other enriched transcripts such as those for cytochrome p450, trypsins, and lipases (Reid et al., 2015). The remodeling of the transcriptome in response to a blood meal likely impacts codon usage as well. In turn, tRNA modifications in females may be necessary to account for the changing codon demand. Anticodon modifications are also sex-specific in mosquitoes and the expression of enzymes involved in anticodon modification is higher in the ovaries than the whole body in *A. aegypti*. Two of the enriched enzymes catalyze the transfer of the modification Q onto G34 of tRNAs. Hypomodified histidine tRNAs with G34 prefer the codon CAC while the presence of Q34 demonstrates no preference and can decode CAC or CAU (Meier et al., 1985). Likewise, in the tobacco mosaic virus (TMV), a hypomodified tyrosine tRNA with G34 suppresses stop codons and fails to halt protein synthesis at the stop codon (Beier and Grimm, 2001; Bienz and Kubli, 1981). On the contrary, the presence of Q34 on another tyrosine tRNA does not act as a suppressor and appropriately terminates translation at the stop codon (Beier and Grimm, 2001; Bienz and Kubli, 1981). In both cases, the presence of Q on the wobble position affects the decoding of codons and subsequently improves translational efficiency. As Q and other anticodon modifications are in higher abundance in females, the enrichment of these enzymes in the ovaries suggests there is potentially sex-specific codon usage that is possibly involved in reproduction.

Some limitations must be addressed to better understand the role of higher tRNA modification in females and their significance in reproduction. Primarily, the method of assessing modification levels is in relative abundance and not absolute abundance. Ideally, the absolute amount of each nucleoside would illustrate the differences between males and females. Additionally, the differences in tRNA modifications detected and their abundances may be caused by differences within the tRNA pool. Previous work in *Anopheles gambiae* notes tRNA transcript abundances are not tissue-specific (Bryant et al., 2020), however, the composition of the tRNA pool may still influence the relative abundance of modifications. For example, if a tRNA transcript containing a specific modification is in generally low abundance, the modification would not be detected. Furthermore, there are discrepancies in the enzyme expression and tRNA modification abundance. In *C. pipiens*, nearly all of the tRNA modification abundances are the same regardless of sex. However, tRNA-modifying enzyme expression is consistently higher in females than in male *C. pipiens*. This demonstrates that the tRNA-modifying enzyme expression is not always reflective of the modification status. This complicated relationship between tRNA-modifying enzyme expression and tRNA modifications in mosquitoes suggests that this may be a fruitful and underexplored area of mosquito biology.

Altogether, we have shown that tRNA modifying enzyme expression and tRNA modification abundance is sex-specific in mosquitoes. The consistent elevation of tRNA modification and enzyme expression suggests that female mosquitoes require more mechanisms for translational efficiency. Modifications can contribute to the thermal stability of tRNAs which may be required in coping with a blood meal. Additionally, the higher levels of anticodon modification in females could suggest preparation for altered codon usage, such as those required to produce materials for location into the eggs. However, future studies to evaluate the effects of a blood meal on tRNA modifications will provide more insight into their relevance concerning blood digestion and reproduction.

## Acknowledgements

Research reported in this publication was partially supported by the National Institute of Allergy and Infectious Diseases of the National Institutes of Health under Award Number R01AI148551. The content is solely the responsibility of the authors and does not necessarily represent the official views of the National Institutes of Health.

